# Identification, optimization, and structural elucidation of chloroacetamide scaffold as covalent inhibitors for Ubiquitin C-terminal Hydrolase L3

**DOI:** 10.64898/2026.05.26.727856

**Authors:** Nagabhushana C. Beeralingappa, Mengyao Lu, Rishi Patel, Nipuni Pannala, Alisha Dhiman, Brittany N. Heil, Ryan D. Imhoff, Emily G. Smith, Mei B. Bahler, Haley L. Marsden, Brittany L. Allen-Petersen, Michael K. Wendt, Chittaranjan Das, Daniel P. Flaherty

## Abstract

The deubiquitinating enzyme, ubiquitin C-terminal hydrolase L3 (UCHL3), has been implicated as a potential therapeutic target for cancer with a role in regulating the DNA damage response pathways. While the target has been studied using genetic methods there is a lack of reliable chemical probes to selectivity target UCHL3. In this study we report hit identification and optimization of a new chemical scaffold that irreversibly inhibits UCHL3. The observed structure-activity relationships are corroborated by ligand-bound crystal structures that confirm covalent adduct formation with the catalytic cysteine of the enzyme. Finally, through gel-shift assays using a ubiquitin activity-based probe we demonstrate on-target engagement with UCHL3 in two cell lines. The work as a whole presents a comprehensive evaluation of the new scaffold that can be utilized to probe UCHL3 in different biological contexts.

## INTRODUCTION

Ubiquitination is a posttranslational modification that covalently binds the signaling protein ubiquitin to target proteins. This modification plays essential roles in several cellular processes, including protein degradation, proteostasis, DNA repair, autophagy, cell signaling, and cell cycle regulation.^1–3^ Ubiquitination can be reversed by deubiquitinating enzymes (DUBs) that remove ubiquitin from proteins and rescue them from degradation.^4^ Cysteine proteases are a primary class of DUBs that catalyze the hydrolysis of the isopeptide bond between the C-terminus of ubiquitin and a lysine side chain on the substrate.^5^ The dysregulation of DUBs are related to multiple health conditions such as cancer, neurodegenerative diseases, cardiovascular diseases, and inflammatory disorders.^6^

Ubiquitin C-terminal Hydrolase L3 (UCHL3) belongs to the UCH family of cysteine proteases. It regulates proteostasis,^5,7,8^ cell cycle control,^9^ and DNA damage repair pathways.^10–12^ UCHL3 promotes the growth of breast cancer,^10,11^ lung cancer,^13^ pancreatic cancer,^7,9^ bladder cancer,^14^ ovarian cancer,^15,16^ and colorectal cancer^17^ via different signaling pathways. One of the common pathways with which UCHL3 is implicated in cancer is through DNA damage repair pathways where deubiquitination and stabilization of RAD51, a key protein in DNA repair, enhances homologous recombination DNA repair and reduces breast cancer cell susceptibility to DNA damage agents such as radiation and chemotherapy.^11^ Knockdown of UCHL3 in triple-negative breast cancer and non-small cell lung cancer models makes tumors cells more susceptible to radiation or topoisomerase inhibitor treatment. Aberrant over expression of UCHL3 is also implicated in regulating aryl hydrocarbon receptor (AhR) stability in triple-negative breast cancer and non-small cell lung cancer.^13,18^ In this case, UCHL3 removes the polyubiquitin chain from AhR and stabilizes AhR in the nucleus, leading to downstream enhanced tumor proliferation. Importantly, genetic depletion of UCHL3 can abrogates this effect.^10,11^ Similar results have been noted in bladder cancer, where UCHL3 activates the Wnt signalling by stabilizing β-catenin^14^ Finally, UCHL3 also deubiquitinates the TNF Receptor Associated Factor 2 (TRAF2) to rescue it from degradation, consequently upregulating the Nuclear Factor kappa-B cells (NF-κB)-associated inflammatory response contributing to the progression of ovarian cancer.^15^ Again, genetic depletion of UCHL3 prevent these tumorigenic events.^7–11,13–18^

The growing evidence of aberrant UCHL3 overexpression as an oncogenic driver suggests its potential as a therapeutic target in cancer therapy. However, the currently available small molecule UCHL3 inhibitors have significant liabilities. The molecule 4,5,6,7-Tetrachloroindan-1,3-dione (TCID) (**Figure 1**.) has been reported as a UCHL3 inhibitor^19^ and although TCID can suppress cancer cell proliferation and downregulate stem cell-like properties in several cancer types,^20^ no structural evidence or target engagement data is available to validate its direct UCHL3 binding in cells. More recently, reports of an AKT inhibitor, perifosine (**Figure 1**), suggested the molecule can enhance the PARP antitumor effect by inhibiting UCHL3-mediated RAD51 activity.^10^ Similarly, the molecule does not inhibit recombinant UCHL3 in standard enzymatic assays and the direct engagement of perifosine and UCHL3 has not been demonstrated.^10,21^ Another strategy employed to probe UCHL3 was development of Ub-variants that selectivity target the enzyme over other DUBs^21,22^, although this approach has clear limitations in only being able to dose whole cells with protein. Taken together, there is a clear need for a new approach to pharmacologically probe UCHL3 to validate its therapeutic potential in cancer.

**Figure 1.**
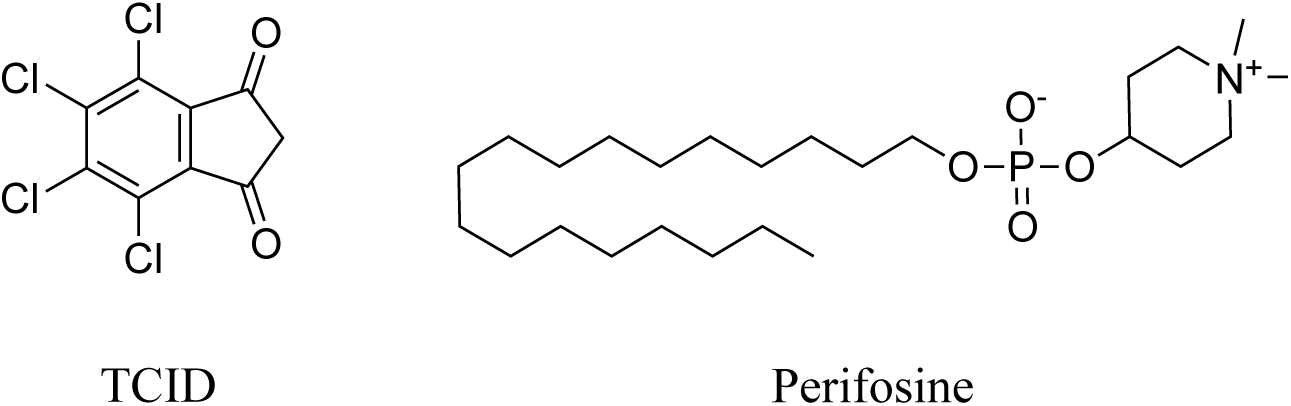
Structures of prior reported UCHL3 inhibitors TCID and Perifosine.

To this end, we conducted a high-throughput screen against UCHL3 to identify hits and tested the hits against the close homolog UCHL1. From this hit identification we carried out structure-activity relationship (SAR) optimization to arrive at a low µM inhibitor. Finally, we report, to the best of our knowledge, the first inhibitor bound UCHL3 crystal structures that were used to gain further insight into engagement with the enzyme.

## RESULTS

### High-throughput screen of covalent fragment library against UCHL3

To identify UCHL3 inhibitors, our team took the approach to exploit the active site cysteine with covalent inhibitors.^23,24^ To achieve this, a 1700-member covalent fragment library from Enamine (Kyiv, Ukraine) containing four known cysteine focused electrophiles; acrylamides, chloroacetamides, activated nitriles and epoxides, was screened against recombinant UCHL3. The screening assay we employed was a fluorescence-based ubiquitin-rhodamine 110 (Ub-Rho) enzymatic activity assay. Prior to screening, a Z’ analysis was conducted resulting in a Z’ of 0.80 providing a sufficient signal-to-noise window to provide reliable data to identify hits from screening. During the screening campaign, iodoacetamide served as a positive control while DMSO served as negative control. A 1-hour preincubation with inhibitor and enzyme was selected based on our previous experience with DUB inhibitors. Fragments were screened in singlet at a concentration of 500 µM with 1-hour preincubation prior to addition of Ub-Rho substrate and subsequent reading. Hits criteria were set at greater than 90% inhibition where percent inhibition was determined compared to DMSO control set to 100% activity. The fragment screen using this hit criteria yielded 34 initial hits (**Figure 2A**). The hits were then validated in triplicate using the Ub-Rho assay resulting in 8 triaged molecules that did not reach 90% inhibition and 26 validated hits for a hit rate of 1.5% (**Figure 2B**). The 26 hits were then counter screened in triplicate against closely related UCHL1 to identify scaffolds selective for UCHL3 (**Figure 2C**). The criteria for the counter screen were <25% percent inhibition against UCHL1. Overall, 14 hits met the hit criteria and were then carried into dose response studies against both DUBs. From the dose response studies, molecule **4** emerged as our prioritized hit due to its potency against UCHL3 (IC_50_ = 57 µM) and selectivity over UCHL1 (IC_50_ >500 µM) (**Figure 2D**). Molecule **4** achieved a ligand efficiency of 0.3 kcal/mol/heavy atom and lipophilic ligand efficiency of 1.44 which are accepted values (LE) or close to accepted (LLE) for fragment hits.

**Figure 2.**
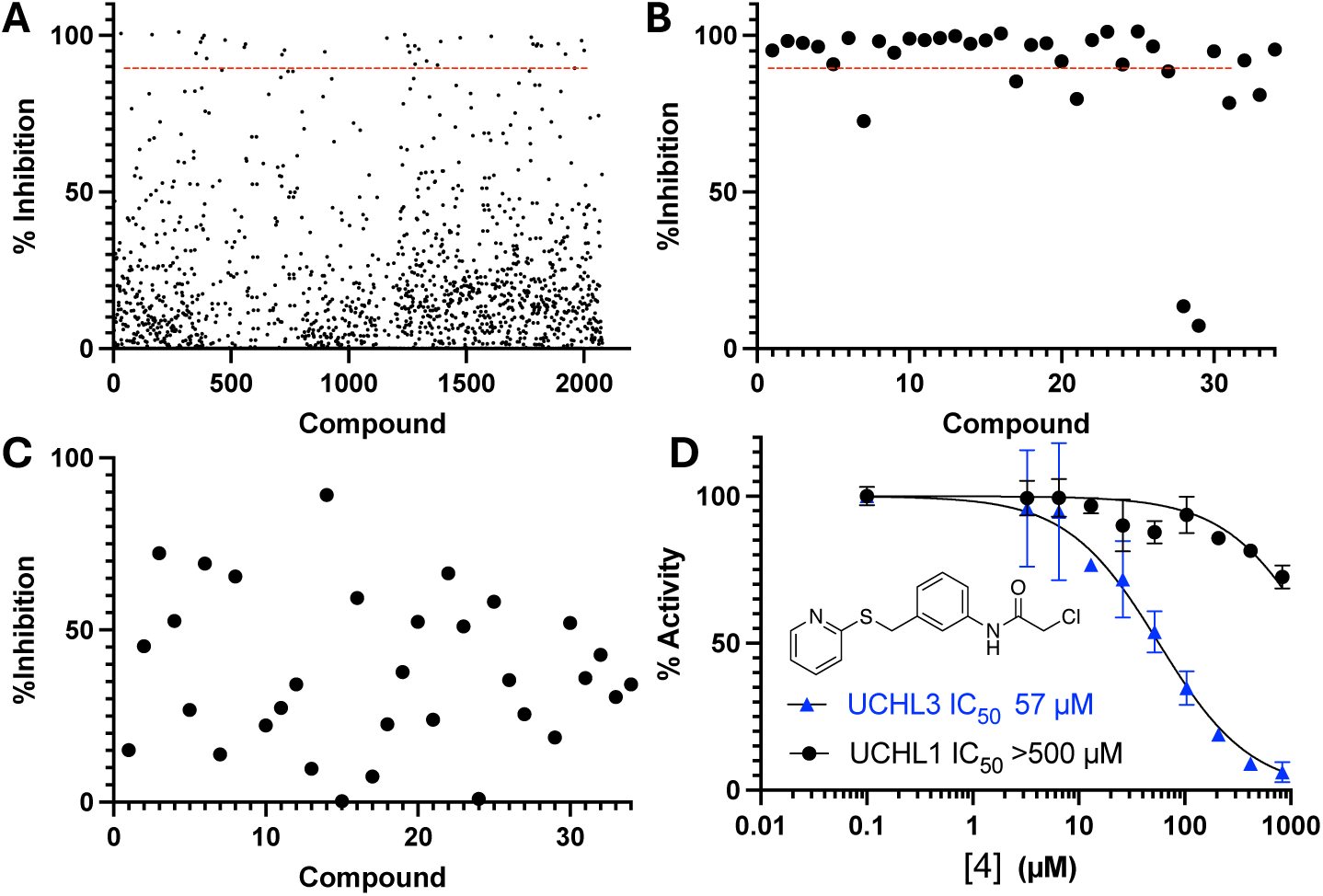
Covalent fragment screening results to identify UCHL3 inhibitors using the ubiquitin rhodamine 110 enzymatic activity assay. (**A**) Primary fragment screen performed in singlet. Red line represents hit cutoff of >90% inhibition. (**B**) Hit validation performed in triplicate. Red line represents hit cutoff of >90% inhibition. (**C**) Counter screen of hit molecules against UCHL1. (**D**) Chemical structure of hit **4** and Ub-Rho 110 assay dose-response.

### Hit molecule **4** covalently modifies catalytic Cys95 of UCHL3

Intact mass spectrometry was conducted to determine the adduct to unmodifies protein ratio. Chloroacetamide electrophiles are relatively reactive toward nucleophilic residues; thus, we wanted to determine if more than one molecule covalently modified UCHL3 at other Cys residues. Hit **4** was incubated with recombinant UCHL3 protein in a 2:1 ligand:protein ratio for 1 hour then evaluated with intact protein mass spectrometry. The data indicated a mass shift of approximately 256 Da (**Table 1**), which equates to a single adduct formation of **4** with UCHL3, presumable at the catalytic Cys95.

**Table 1.**
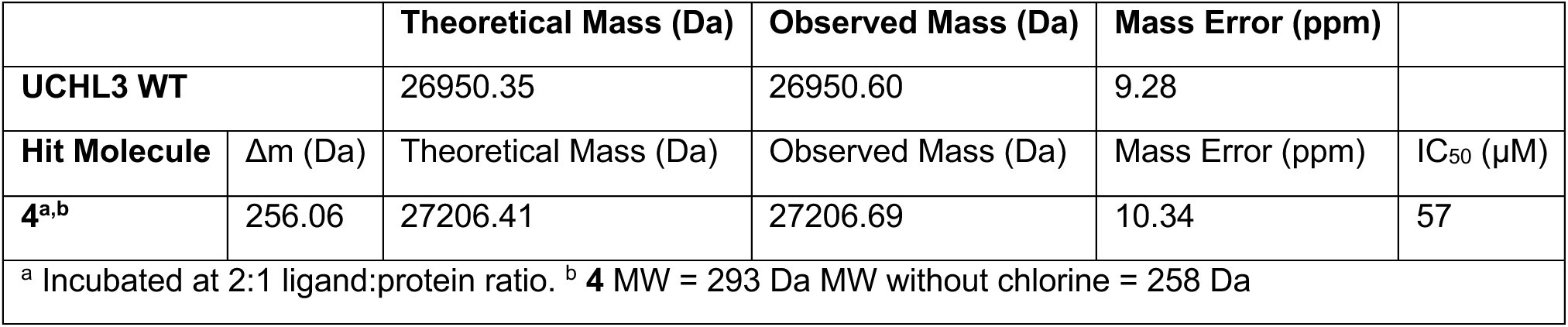
Intact Protein Mass Spectrometry for UCHL3 with and without 4.

| **Table 1.** Intact Protein Mass Spectrometry for UCHL3 with and without **4** |  |  |  |  |  |
| --- | --- | --- | --- | --- | --- |
|  |  | **Theoretical Mass (Da)** | **Observed Mass (Da)** | **Mass Error (ppm)** |  |
| **UCHL3 WT** |  | 26950.35 | 26950.60 | 9.28 |  |
| **Hit Molecule** | $\Delta m$ (Da) | Theoretical Mass (Da) | Observed Mass (Da) | Mass Error (ppm) | $IC_{50}$ ($\mu M$) |
| **4<sup>a,b</sup>** | 256.06 | 27206.41 | 27206.69 | 10.34 | 57 |
| <sup>a</sup> Incubated at 2:1 ligand:protein ratio. <sup>b</sup> **4** MW = 293 Da MW without chlorine = 258 Da |  |  |  |  |  |

The hit **4** was carried forward into crystallographic studies to determine binding pose with UCHL3. The molecule was pre-incubated with recombinant UCHL3 (1:5 ratio) for 1 hour at room temperature to form the complex, as indicated by the appropriate mass shift (**Table 1b**), and subsequently purified and crystallization trials were conducted to determine its pose. We solved the X-ray crystal structure of UCHL3 covalently bound to **4** to a resolution of 1.95 Å in the P2_1_ space group (ID: pdb_000013CU, **Figure 3**, data collection and refinement statistics presented in **Table S1**). This structure showed three chains in the asymmetric unit with a single ligand covalently bounds to the catalytic Cys95 on chains A and B but no ligand was observed in Chain C, corroborating our observation of partial inhibition from intact mass spectrometry. The poses for molecule **4** in chains A and B were found to be at the interface of those chains (**Figure 3A**). Each chain viewed individually demonstrates the flexibility of **4**, particularly the pyridine-thioether tail, rotating in different directions (**Figure 3B – D**). The presence of two poses is likely attributed to the expected low affinity of the ligand for the target at the hit identification stage, making it difficult to select one specific pose that may be more relevant for further ligand optimization. Therefore, the structures were not utilized for design of analogs. Nevertheless, they confirmed single-adduct formation of **4** to the catalytic Cys95 in the covalent adduct.

**Figure 3.**
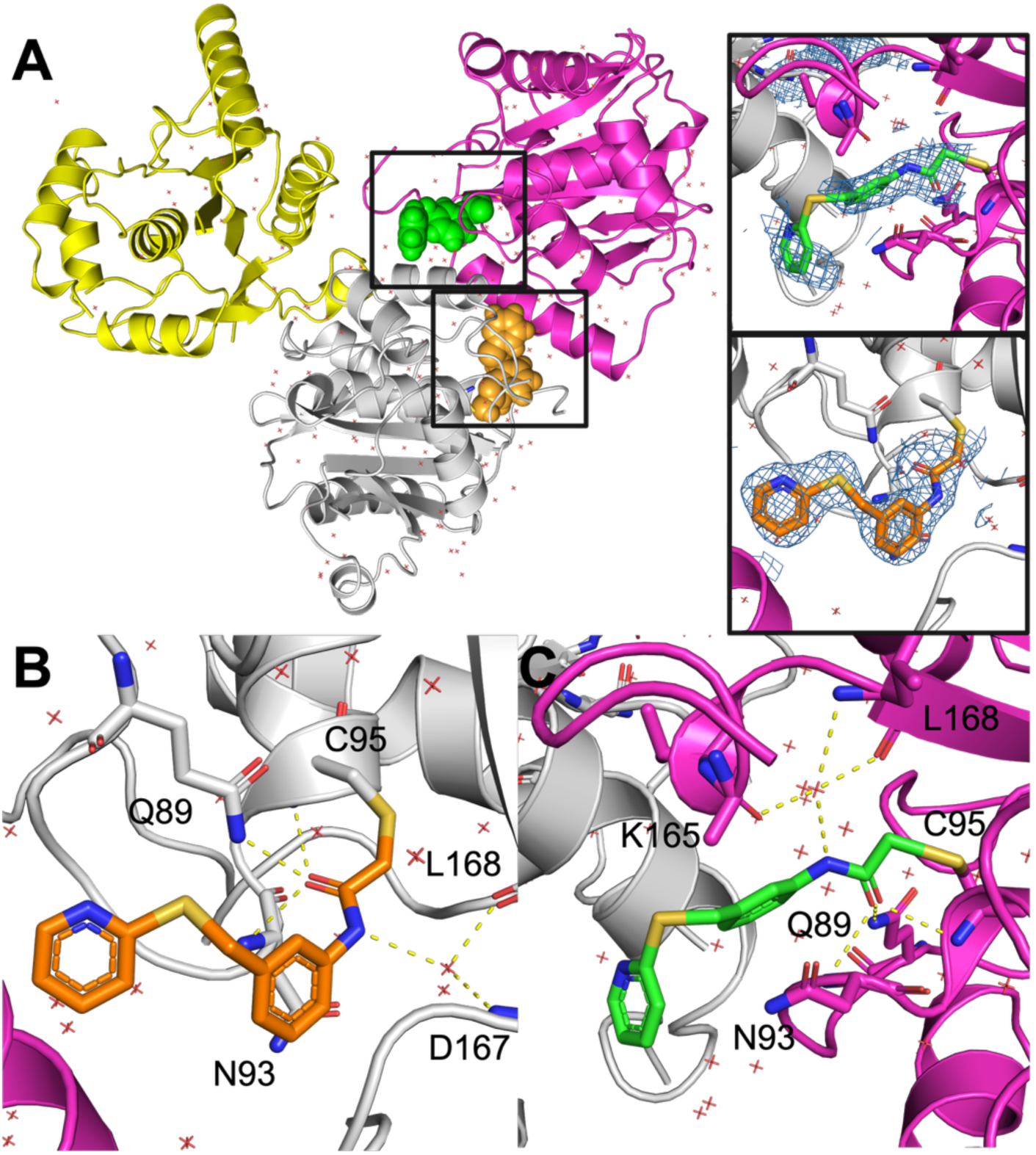
Ligand-bound crystal structure of UCHL3 in complex with **4** at 1.95 Å (ID: pdb_000013CU). (**A**) Full asymmetric unit showing three UCHL3 chains with **4** depicted as spheres in each chain. Omit |2*F*_0_ – *F*_c_| electron density map for each chain shown in insets. (**B**) Chain A (gray) with **4** (orange sticks). (**C**) Chain B (magenta) with **4** (green sticks). Water molecules as red tetrahedrons and hydrogen bonds depicted by yellow dashed lines.

### Chemistry

Based on the structural architecture of chloroacetamides, they have been synthesized by using different reaction strategies (**Schemes 1** – **5**). **Scheme 1** depicts the synthetic strategy for compounds **4 – 10** using commercially available 3-nitrobenzylbromide **1**. The desired thiophenol/phenol/Boc-amines were made to react with 3-nitrobenzylbromide **1** under potassium carbonate conditions at 60 °C generated corresponding nitro derivatives (**2a-g**). Iron-mediated reduction of nitro to amine followed by the treatment with desired acid chlorides generated compounds **4-9**. Finally, boc-deprotection of intermediate **10a** with HCl afforded the hydrochloride salt of compound **10**. Similarly, commercially available amines **11** have been subjected to react with chloroacetyl chloride in the presence of Et_3_N, giving corresponding chloroacetamides **12-17** (**Scheme 2**).

**Scheme 1:**
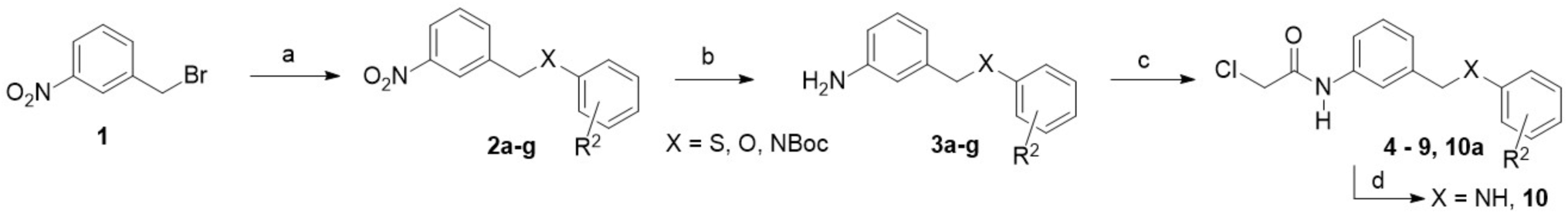
Reagents and reaction conditions: (a) Thiol/phenol/amine(pyridine derivatives as well) (1 Eq), K_2_CO_3_ (1.2 Eq), DMF, 60 _°_C, 59 -89%; (b) Fe powder (5 eq), 1:2:2 H_2_O, ethanol, acetic acid, sonication, RT, 45 -92%; (c) Acid chloride (1.1 Eq), Et_3_N (1.5 Eq), THF, 0 _°_C – RT, 40 -66%; (d) HCl in Et_2_O, Et_2_O, 0 _°_C – RT, 77%.

**Scheme 2:**
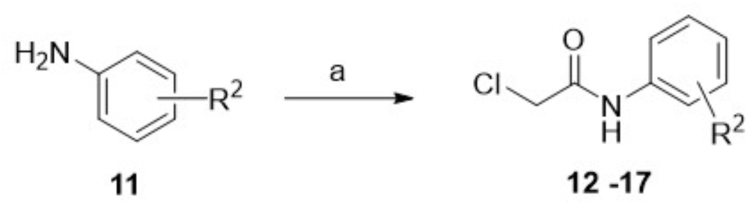
Reagents and reaction conditions: (a) chloroacetyl chloride (1.1 Eq), Et_3_N (1.5 Eq), THF, 0 _°_C – RT, 70 -90%.

**Scheme 3:**
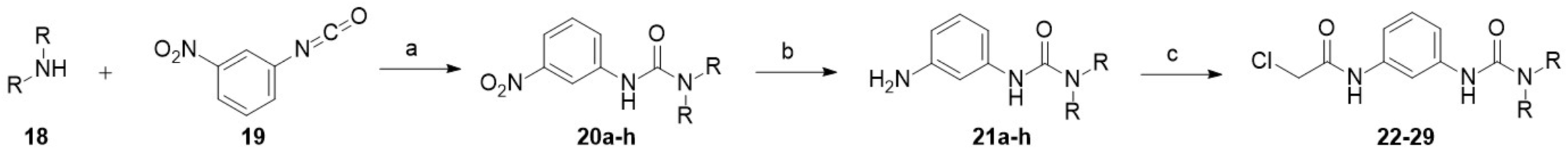
Reagents and reaction conditions: a) CH_3_CN, RT, 2 hrs, 82 – 92%; (b) Fe powder (5 eq), 1:2:2 H_2_O, ethanol, acetic acid, sonication, rt, 2 hr, 50 -67 %; (c) chloroacetyl chloride (1.1 Eq), Et_3_N (1.5 Eq), THF, 0 _°_C – RT, o/n, 27 -82%.

**Scheme 4:**
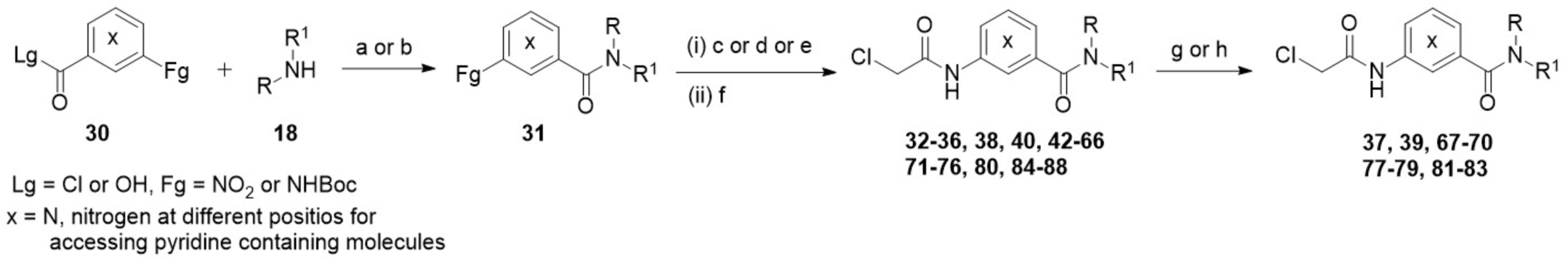
Reagents and reaction conditions: (a) With acid chlorides: Et_3_N (1.5 Eq), THF, 0 ^°^C – RT, 60 -85%; (b) With Acids: HATU (1.2 Eq), DIPEA (2.2 Eq), DMF, RT, 50 -90%; (c) Fe (5 Eq), NH_4_Cl (5 Eq), THF:MeOH:H_2_O, 60 ^°^C; or (d) H_2_, Pd/C, MeOH, RT (if no benzyloxy groups present) or (e) For Boc deprotection: TFA (15 Eq), DCM, 0 ^°^C – RT; (f) Chloroacetyl chloride (1.1 Eq), Et_3_N (1.5 Eq), 0 ^°^C – RT, 40 -80%; (g) H_2_, Pd/C (10% w/w), MeOH, RT (For deprotection of benzyloxy groups) or (h) HCl in Et_2_O, Et_2_O, 0 ^°^C – RT (For Boc deprotection **(41 & 79)** – HCl salts are isolated).

**Scheme 5:**
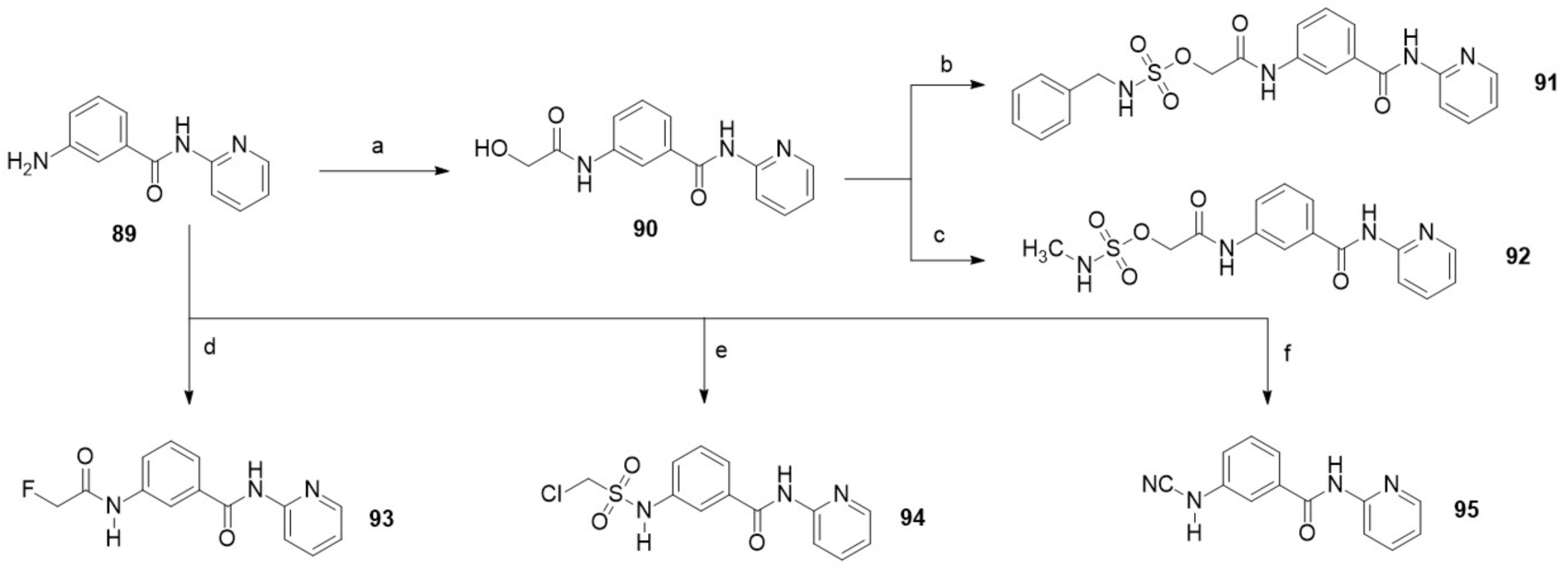
Reagents and reaction conditions: a) 2-hydroxyacetic acid (1.3 Eq), EDCI (1.3 Eq), HOBt (1.3 Eq), DIPEA (1.6 Eq), DCM, RT, 12 hrs, 12%; b) benzylsulfamoyl chloride (1.2 Eq), DIPEA (1.2 Eq), DCM, 0 ^°^C – RT; 60 min, 68%; c) methylsulfamoyl chloride (1.2 Eq), DIPEA (1.2 Eq), DCM, 0 ^°^C – RT, 60 min, 30%; d) Fluoroacetyl chloride (1.1 Eq), Et_3_N (1.3 Eq), DCM, 0 ^°^C – RT, 3 hrs, 78%; e) chloromethanesulfonyl chloride(1.1 Eq), Et_3_N (1.3 Eq), DCM, 0 ^°^C – RT, 4 hrs, 52%; f) Zn(CN)_2_ (1.0 Eq), N-chlorosuccinimide (1.5 Eq), CH_3_CN:H_2_O, 0 ^°^C – RT, 16 hrs, 80%.

Compounds **22-29** have been synthesized by following a three-step procedure using commercially available starting materials such as 3-nitrophenylisocyanate and desired amines (**Scheme 3**). The reaction between amine **18** and 3-nitrophenylisocyanate **19** in acetonitrile provided corresponding urea derivatives **20a-h**. Reduction of nitro to amine under iron/AcOH conditions generated amines **21a-h**. Finally, amines **21a-h** have been treated with chloroacetyl chloride in the presence of Et_3_N to get corresponding chloroacetamides **22-29**.

Most of the chloroacetamides **32 – 88** were synthesized by using general reaction strategy as shown in **Scheme 4**. This strategy has been designed based on substrate availability and functional group tolerance. Commercially available amines **18** were made to react either with acid chloride or acid **30** under Et_3_N or HATU/DIPEA to get corresponding amides. The desired amines were accessed either by nitro reduction using Fe/NH4Cl or H_2_-Pd/C reaction conditions or by Boc-deprotection using TFA. The crude amines were acylated with chloroacetyl chloride to generate compounds **32 – 36** and **38**. Finally, some of the intermediates were converted to desired compounds **37, 39, 67-70, 77-79, 81-83** either by using palladium-catalyzed hydrogenation or by Boc-deprotection with TFA. To have different warheads in place of chloroacetamides, the different reaction strategy was followed as shown in **Scheme 5**. The amine **89** was coupled with 2-hydroxyacetic acid under HATU/DIPEA conditions, affording alcohol **90**, which was further reacted with benzylsulfamoyl and methylsulfamoyl chloride to get corresponding sulfamides **91** and **92** respectively. Similarly, compound **89** was treated with fluoroacetyl chloride and chloromethanesulfonyl chloride to access corresponding amides **93** and **94**. Amine **89** on treatment with zinc cyanide and *N*-chlorosuccinimide generated *N*-cyano derivative **95.**

### Structure-activity relationship of cysteine targeted irreversible inhibitors of UCHL3

The inhibitory activities of target compounds against UCHL3 were assessed using the standard fluorescence-based ubiquitin-rhodamine 110 (Ub-Rho) enzymatic activity assay, a well-established method for evaluating the catalytic activity assays, inhibitor screening and profiling of DUBs. The enzymatic inhibitory potency of tested chloroacetamides and other cysteine-focused electrophilic derivatives was displayed by IC_50_ values illustrated in **Tables 1-5**.

**Table 1:**
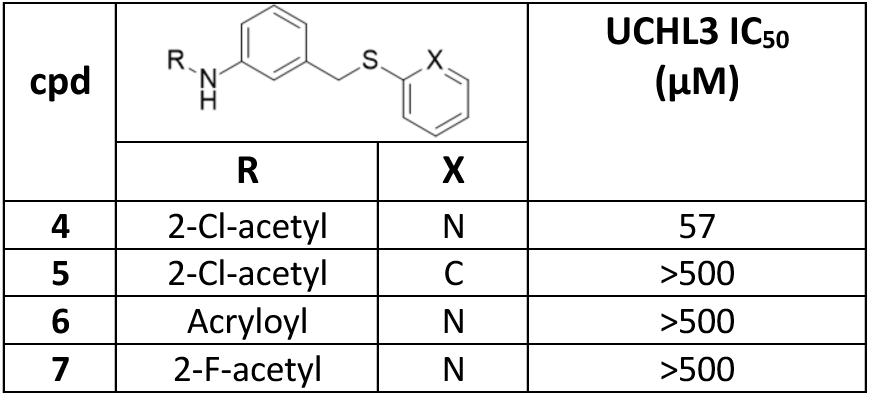
UCHL3 activity for compounds 4-7.

| **Table 1: UCHL3 activity for compounds 4-7** |  |  |  |
| --- | --- | --- | --- |
| cpd |  |  | UCHL3 IC<sub>50</sub><br>(μM) |
|  | R | X |  |
| **4** | 2-Cl-acetyl | N | 57 |
| **5** | 2-Cl-acetyl | C | >500 |
| **6** | Acryloyl | N | >500 |
| **7** | 2-F-acetyl | N | >500 |

First, initial targeted modifications to the pendant pyridine and electrophile were evaluated. Modification of hit molecule **4** either by changing pyridine with a phenyl ring or by changing to other electrophiles (**5** – **7**; **Table 1**) were all inactive towards UCHL3. This revealed that chloroacetamide motif is necessary to maintain the proper electrophilic character and positioning for nucleophilic attack by the catalytic cysteine (Cys95).

The next cohort of analogs maintained the chloroacetamide electrophile while exploring various aspects of the pharmacophore. Analogs were designed to evaluate alternative linkers, changes to the pendant pyridine, and probed connectivity and makeup of the central phenyl core (**Table 2**). Swapping the thioether to either an ether or amine linker (**8** – **10**) were found to reduce UCHL3 activity compared to **4.** Additional ether linker/core modification also resulted in lost activity versus UCHL3, except for ether linker combined with a pendant phenyl in **12** which improved potency (IC_50_ = 24 µM). Inserting a pyridine in the central core for compound **15** was tolerated with IC_50_ = 47 µM.

**Table 2:** UCHL3 activity for compounds 8-47.

| **Table 2: UCHL3 activity for compounds 8-47** |  |  |  |  |  |  |  |  |  |
| --- | --- | --- | --- | --- | --- | --- | --- | --- | --- |
| Where X = N, pyridines |  |  |  |  |  |  |  |  |  |
| cpd | Y | R | X | UCHL3<br>$IC_{50}$ ($\mu M$) | cpd | Y | R | X | UCHL3<br>$IC_{50}$ ($\mu M$) |
| **8** | 3-CH<sub>2</sub>-O- | 2-pyridyl | C | 74 | **32** | 3-NHCOCH<sub>2</sub>- | Ph | C | 20 |
| **9** | 3-CH<sub>2</sub>-O- | 2-naphthyl | C | >500 | **33** | 3-CH<sub>2</sub>CONH- | Ph | C | 93 |
| **10** | 3-CH<sub>2</sub>-NH- | 2-pyridyl | C | 118 | **34** | 3-CONHCH<sub>2</sub>- | 2-pyridyl | C | 243 |
| **12** | 3-O- | Ph | C | 24 | **35** | 3-NHCOCH<sub>2</sub>- | 2-pyridyl | C | 251 |
| **13** | 4-O- | Me | 3-N | >500 | **36** | 3-CONHCH<sub>2</sub>- | 3-OBn-Ph | C | 96 |
| **14** | 4-O- | Ph | 3-N | >500 | **37** | 3-CONHCH<sub>2</sub>- | 3-OH-Ph | C | 78 |
| **15** | 3-O- | Ph | 5-N | 47 | **38** | 3-NHCOCH<sub>2</sub>- | 3-OBn-Ph | C | 68 |
| **16** | 3-SO<sub>2</sub>NH- | Bn | C | 396 | **39** | 3-NHCOCH<sub>2</sub>- | 3-OH-Ph | C | 57 |
| **17** | 3-NHSO<sub>2</sub>- | Bn | C | 370 | **40** | 3-CONH- | Ph | C | 75 |
| **22** | 3-NHCON(Me)- | Ph | C | 88 | **41** | 3-CONH- | 4-piperidyl | C | >500 |
| **23** | 3-NHCONH- | Ph | C | 27 | **42** | 3-NHCO- | 2-pyridyl | C | 32 |
| **24** | 3-NHCONH- | 2-pyridyl | C | 37 | **43** | 3-CONH- | 2-pyridyl | C | 13 |
| **25** | 3-NHCON(Me)- | Ph | C | 88 | **44** | 3-CONH- | 3-pyridyl | C | 45 |
| **26** | 3-NHCONH- | Bn | C | 29 | **45** | 3-CONH- | 2-pyridyl | 5-N | 89 |
| **27** | 3-NHCONH- | 4-Cl | C | >500 | **46** | 3-CONH- | 2-pyridyl | 4-N | >500 |
| **28** | 3-NHCONH- | Cy | C | 29 | **47** | 3-CONH- | 2-pyridyl | 2-N | >500 |
| **29** | 4-NHCONH- | Ph | C | >500 |  |  |  |  |  |

We then evaluated the inhibitory activity of compounds by employing more polar and less flexible linker options. These included sulfonamide-, urea-, and amide-based linkers. Sulfonamide containing molecules were found to be ineffective (**16** and **17**), whereas urea derivatives (**22** – **29**) were generally well tolerated with three (**23**, **26**, and **28**) displaying improved UCHL3 activity (IC_50_ values 27 -29 µM) compared to that of parent molecule **4**. Notably, swapping the urea nitrogen for a methylene group near the pendant phenyl in compound **32** (Y = 3-NHCOCH_2_-, R = Ph) exhibited another step-up in potency (IC_50_ = 20 µM) whereas reversing the orientation of this extended amide linker (**33**; Y = 3-CH_2_CONH-) was 4-fold weaker in terms of UCHL3 activity. This trend was explored with other pendant groups (**34** – **39**) but none were as effective at analog **32**.

Next, we shortened the amide linker by removing the methylene (**40** – **47**, **Table 2**). This was detrimental to activity in the case of analog **40**, with a pendant phenyl group combined with the amide linker was 3 – 4-fold less active than the matched molecular pair with the methylene inserted. However, when the amide linker was combined with 2-pyridine pendant group another boost in activity was observed. In this case, the 2-pyridine containing **42** (Y = 3-NHCO-) showed moderate inhibition with IC_50_ = 32 µM while its positional isomer **43** (Y = 3-CONH-) exhibited the most potent inhibitory activity against UCHL3 observed thus far in the study with IC_50_ = 13 µM. Finally, we maintained the amide linker/orientation and 2-pyridine pendant group and again probed the necessity of central aromatic ring. Any modification to the central phenyl core, whether it be insertion of pyridine (**45** – **47**, **Table 2**) or a cyclic alkane core (**48** – **51**; **Table 3**), all resulted in reduced or full loss of activity against UCHL3. The SAR from this set demonstrated the amide linker was among the most well tolerated and yielded the most potent analog when combined with a 2-pyridine pendant group. With this information we next evaluated the central phenyl ring.

**Table 3:**
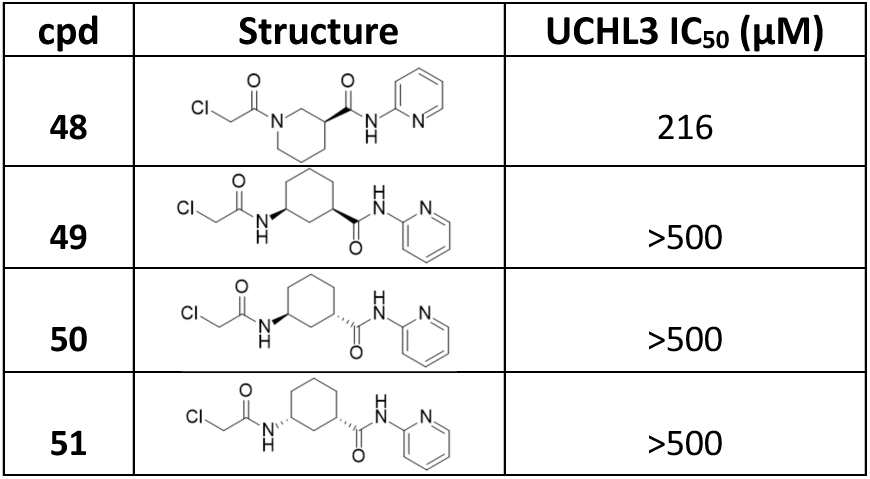
UCHL3 activity for compounds 48-51.

| Table 3: UCHL3 activity for compounds 48-51 |  |  |
| --- | --- | --- |
| cpd | Structure | UCHL3 $IC_{50}$ ($\mu M$) |
| **48** |  | 216 |
| **49** |  | >500 |
| **50** |  | >500 |
| **51** |  | >500 |

After confirming the requirement of the chloroacetamide motif and central phenyl ring system and preference for the amide linker we sought to expand chemical space on the pendant group and evaluate UCHL3 activity (**Table 4**). Many electron-donating and electron-withdrawing groups were evaluated with some observed trends. Among the methyl substituted analogs, placement of CH_3_ at the 2- or 3-positions (**52** and **53**, IC_50_ of 34 and 19 µM, respectively) were tolerated while the 4-methyl was not (**54**). A similar trend was observed with methoxy substitution (**59** – **61**) and dimethylaniline (**62** – **64**) with 4-position not being tolerated. Fluorine at any position was not tolerated (**55** – **58**).

**Table 4:** UCHL3 activity for compounds 52-88 against UCHL3.

| Table 4: UCHL3 activity for compounds 52-88 against UCHL3 |  |  |  |  |  |  |  |  |  |  |  |
| --- | --- | --- | --- | --- | --- | --- | --- | --- | --- | --- | --- |
| cpd | R | R<sup>1</sup> | R<sup>2</sup> | R<sup>3</sup> | UCHL3 IC<sub>50</sub> (μM) | cpd | R | R<sup>1</sup> | R<sup>2</sup> | R<sup>3</sup> | UCHL3 IC<sub>50</sub> (μM) |
| **52** | 2-Me-Ph | H | H | H | 34 | **71** | 6-Me-2-pyridyl | H | H | H | 29 |
| **53** | 3-Me-Ph | H | H | H | 19 | **72** | 4-Me-2-pyridyl | H | H | H | 27 |
| **54** | 4-Me-Ph | H | H | H | >500 | **73** | 6-OMe-2-pyridyl | H | H | H | 19 |
| **55** | 2-F-Ph | H | H | H | >500 | **74** | 4-OMe-2-pyridyl | H | H | H | 16 |
| **56** | 3-F-Ph | H | H | H | >500 | **75** | 3,5-di-OMe-Ph | H | H | H | 49 |
| **57** | 4-F-Ph | H | H | H | 338 | **76** | 4,6-di-OMe-2-pyridyl | H | H | H | 80 |
| **58** | 6-F-2-pyridyl | H | H | H | >500 | **77** | 2-OH-5-Me-Ph | H | H | H | 24 |
| **59** | 2-OMe-Ph | H | H | H | 30 | **78** | 2-OMe-5-Me-Ph | H | H | H | 34 |
| **60** | 3-OMe-Ph | H | H | H | 19 | **79** | 2-NH<sub>2</sub>-Ph | H | H | H | 45 |
| **61** | 4-OMe-Ph | H | H | H | >500 | **80** | 2-acetamido-Ph | H | H | H | 134 |
| **62** | 2-NMe<sub>2</sub>-Ph | H | H | H | 26 | **81** | 2-CH<sub>2</sub>OH-Ph | H | H | H | 75 |
| **63** | 3-NMe<sub>2</sub>-Ph | H | H | H | 98 | **82** | (1*R*,2*R*)-2-OH-Cy | H | H | H | 53 |
| **64** | 4-NMe<sub>2</sub>-Ph | H | H | H | 463 | **83** | (1*S*,2*R*)-2-OH-Cy | H | H | H | >500 |
| **65** | 2-pyridyl | H | Me | H | >500 | **84** | 2-OBn-Ph | H | H | H | >500 |
| **66** | 2-pyridyl | Me | H | H | >500 | **85** | 3-OBn-Ph | H | H | H | >500 |
| **67** | 2-pyridyl | H | H | OH | >500 | **86** | 2-OBn-5-Me-Ph | H | H | H | >500 |
| **68** | 2-OH-Ph | H | H | H | 14 | **87** | 2-OBn-5-OMe-Ph | H | H | H | >500 |
| **69** | 3-OH-Ph | H | H | H | 15 | **88** | (1*R*,2*R*)-2-OBn-Cy | H | H | H | >500 |
| **70** | 4-OH-Ph | H | H | H | 20 |  |  |  |  |  |  |

Next, we probed the role of the amide hydrogen by introduction of a methyl group in each of the amide motifs (**65** and **66**) which resulted in loss of UCHL3 activity suggesting the N-H is critical to target engagement. Additionally, applying a hydroxy group in the central ring was also found to be unfavorable to target inhibition (**67**). Notably, substitution with hydroxy groups on the pendant phenyl was well tolerated with the 2- and 3-OH derivatives, **68** and **69**, exhibiting comparable potency to the top analog **43** at IC_50_ values of 14 µM and 15 µM, respectively. The next cohort of analogs explored combinations of pendant modifications, including pairing methyl and methoxy substituents with the pyridine heterocycle (**71** – **81**). Most of the analogs were found to be reasonably effective in inhibiting the UCHL3 activity and among the group analog **74** exhibited the strongest potency with IC_50_ = 16 µM. The final group of nearest neighbor analogs explore the structural diversity around chiral cyclohexanol derivatives and substituted benzene (**82** – **88)** but were found to be inactive towards UCHL3 except for **82** with an IC_50_ value of 53 µM.

Once we identified a more potent scaffold in **43** we revisited the activity of different warheads combined with the core structure of **43** (**Table 5**). Sulfamate electrophiles have been demonstrated to react similarly to chloroacetamides and maintain improved stability in cells and *in vivo*;^25^ therefore, we evaluated the benzenesulfame analog **91** and methylsulfamate **92**. Among these, only **91** exhibited any UCHL3 activity at 70 µM, though much reduced compared to **43**. Similarly, we observed no activity with the incorporation of a fluoroacetamide warhead (**93**), similar to the fluoromethylketone warhead that has previously demonstrated activity versus closely related UCHL1.^26,27^ (**91-93**). Finally, neither (chloromethyl)sulfonamido) nor cyano warheads potent activity versus UCHL3 (**94-95**).

**Table 5:**
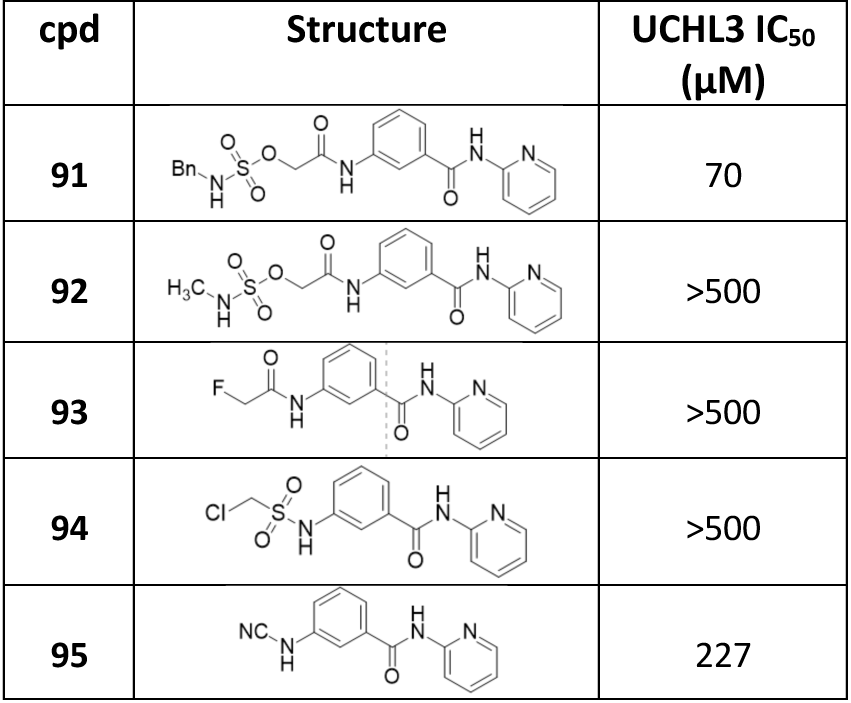
UCHL3 ac/vity for compounds 91-95.

| **Table 5: UCHL3 activity for compounds 91-95** |  |  |
| --- | --- | --- |
| **cpd** | **Structure** | **UCHL3 IC<sub>50</sub> (μM)** |
| **91** |  | 70 |
| **92** |  | >500 |
| **93** |  | >500 |
| **94** |  | >500 |
| **95** |  | 227 |

Once SAR was complete against UCHL3, we selected inhibitors with IC_50_ < 20 µM against UCHL3 to test their activity against its close homolog UCHL1 (**Table 6**). In general, almost all analogs were not active against UCHL1 with two exceptions. Compound **32** exhibited measurable, yet weak, inhibitory activity against UCHL1, with IC_50_ = 313 µM. Compound **69** showed moderate inhibition against UCHL1(IC_50_ = 45 µM), with a 3-fold selectivity for UCHL3. Interestingly, the top-performing analogs in terms of UCHL3 activity all exhibited no UCHL1 inhibition at 500 µM, indicating at least 30-fold selectivity for UCHL3.

**Table 6:** UCHL1 activity for L3-potent compounds.

| Table 6: UCHL1 activity for L3-potent compounds |  |  |  |
| --- | --- | --- | --- |
| cpd | IC<sub>50</sub> (μM) | cpd | IC<sub>50</sub> (μM) |
| 32 | 313 | 69 | 45 |
| 43 | >500 | 70 | >500 |
| 53 | >500 | 73 | >500 |
| 60 | >500 | 74 | >500 |
| 68 | >500 |  |  |

### Kinetics of irreversible inhibition of UCHL3

The top-performing covalent analogs were further evaluated by measuring *k*_inact_/K_I_ using a reported protocol.^28^ The *k*_inact_ is the rate constant for covalent bond formation between the inhibitor and the enzyme under saturating conditions, and K_I_ is the constant that reflects the inhibitor’s affinity for the enzyme before the covalent interaction. The observed rate constant (*k*_obs_) was determined from an enzymatic activity assay of UCHL3 at different inhibitor concentrations across different preincubation times of the inhibitor and enzyme. The *k*_obs_ values were plotted as a function of increasing inhibitor concentration, generating a curve for **68** that could be fit using the equation Y = *k*_inact_*X/(K_I_ + X) (**Figure 4**). These values were determined to be K_I_ = 274.7 µM and *k*_inact_ = 0.005831 s^−1^, resulting in a *k*_inact_/K_I_ value of 21.23 M^−1^ s^−1.^ For **43** and **69**, the *k*_obs_ were fitted linearly against increasing inhibitor concentrations, and the slopes of the linear regressions yield the ratio of *k*_inact_/K_I_ together but not the individual parameters (**Figure 4**). The *k*_inact_/K_I_ values for **43** and **69** were determined to be 11.57 M^−1^ s^−1^ and 14.74 M^−1^ s^−1^, respectively. The difference in data analysis method between **68** and the other molecules was due to **68** having a better affinity for forming the reversible enzyme-inhibitor complex with UCHL3, while **43** and **69** bind relatively slowly to the enzyme but rely more on the inactivation step for inhibition. These results indicate that molecules, while inhibitory, still have room to improve for affinity to UCHL3.

**Figure 4.**
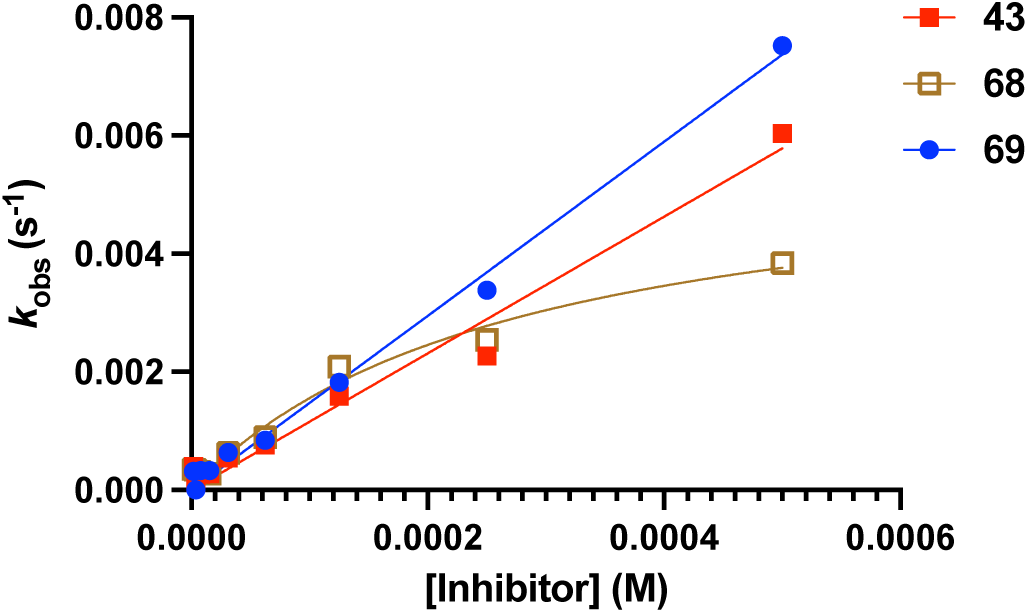
Inactivation kinetics for top-performing analogs.

### Structural elucidation of binding for UCHL3 inhibitors

We next set out to elucidate the interactions of inhibitors bound to UCHL3 using X-ray crystallography similarly as described above for the hit molecule **4**. We were able to resolve high-resolution ligand-bound structures for analogs **43** (ID: pdb_000013CV, 1.95 Å), **53** (ID: pdb_000013CW, 2.48 Å), **60** (ID: pdb_000013DF, 1.84 Å), and **69** (ID: pdb_000035YH, 2.60 Å) bound to UCHL3 (**Figure 5**, ligand density in **Figure S1** and crystallographic data statistics in **Table S1** – **S3**). The structures revealed four subunits with a single ligand bound to the Cys95 for each subunit and resided at the interface between chains. All inhibitors maintain interactions between the acetamide carbonyl oxygen with the oxyanion hole of the enzyme. The oxyanion hole in the UCH DUB family consist of amide functional groups, either on a glutamine side-chain or a backbone amides, where the nitrogen atoms form hydrogen bonds with the substrate carbonyl oxygen of the scissile peptide bond to stabilize the transition state thereby facilitating the catalytic turnover to cleave ubiquitin.^29,30^ This placement of the carbonyl into the oxyanion hole positions the amide nitrogen of the acetamide toward the backbone carbonyl of Leu168 for all three ligand-bound structures. These experimentally observed interactions between the inhibitors and UCHL3 corroborate the observed SAR, particularly with respect to the placement of the carbonyl with respect to the electrophilic carbon. The carbonyl placement in the oxyanion hole likely positions the electrophilic center in the proper orientation for nucleophilic attack by Cys95 and explains why alternative warheads are not useful for this scaffold against UCHL3. Moreover, methylation of the chloroacetamide nitrogen (**66**) abrogated activity when comparing to the matched molecular paired analog **43**, likely due to disruption of the interaction with the backbone carbonyl of Leu168 leading to altering the position of the warhead for efficient reactivity.

**Figure 5.**
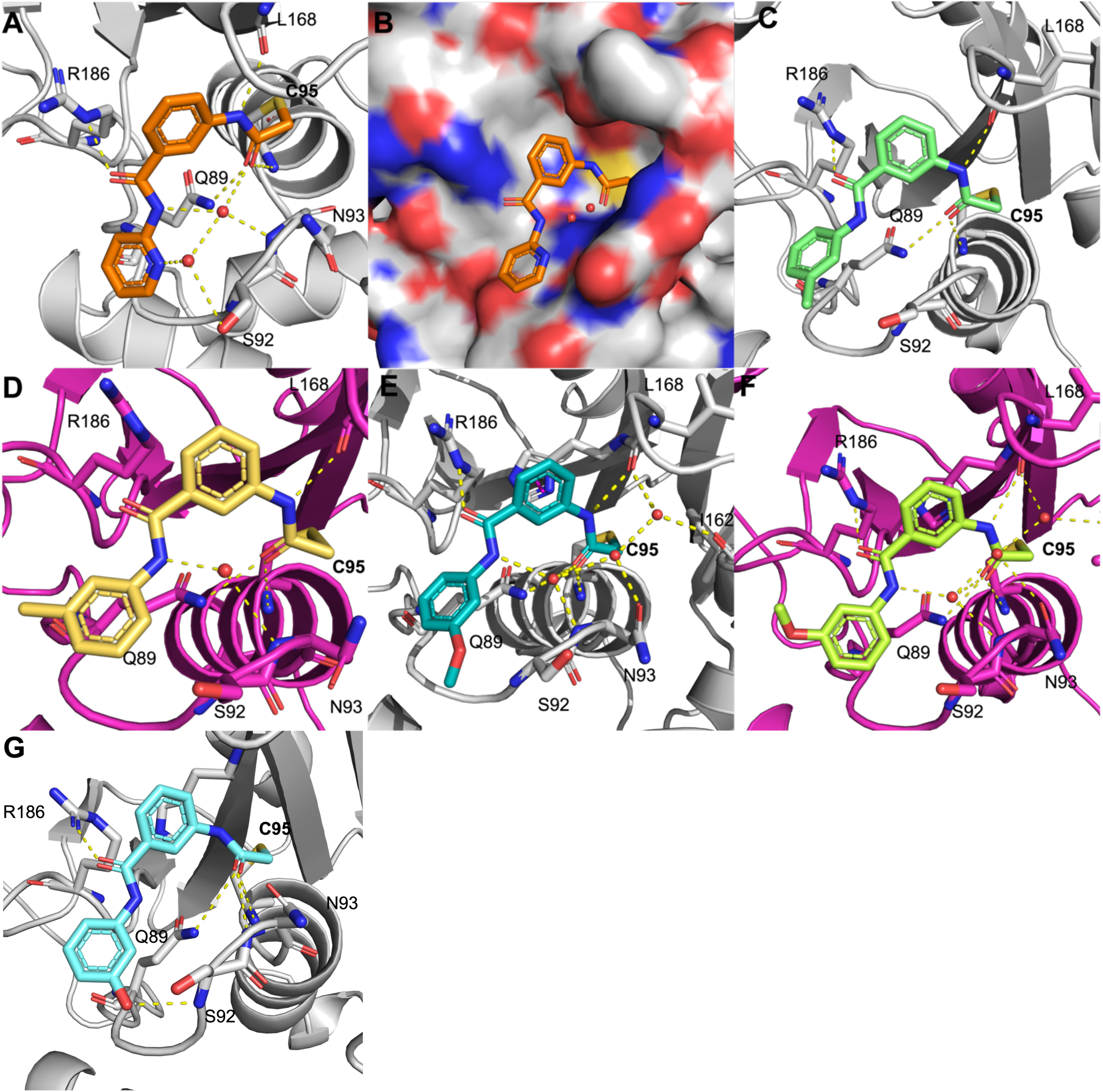
Ligand-bound structures of covalent inhibitors bound to UCHL3. In all panels, water molecules depicted as red spheres and hydrogen bonds depicted by yellow dashed lines. Residues involved in ligand binding shown as sticks and labeled. (**A**) **43** (orange sticks) bound to UCHL3 (chain A; gray cartoon; ID: pdb_000013CV). (**B**) **43** bound to UCHL3 (chain A; gray surface) in surface representation. (**C**) **53** (mint sticks) bound to UCHL3 (chain A; gray cartoon; ID: pdb_000013CW). (**D**) **53** (gold sticks) bound to UCHL3 (chain B; magenta cartoon;). (**E**) **60** (teal sticks) bound to UCHL3 (chain A; gray cartoon; ID: pdb_000013DF). (**F**) **60** (light green sticks) bound to UCHL3 (chain B; magenta cartoon). (**G**) **69** (sky blue sticks) bound to UCHL3 (chain A; gray cartoon; ID: pdb_000035YH).

The importance of the amide linker position and connectivity between the central aromatic core and the pendent group was also illuminated by the ligand-bound structures. For example, the placement of the amide carbonyl in the linker on all analogs consistently formed a hydrogen bond with the side chain of Arg186 of UCHL3 (**Figure 5**). This pulled the carbonyl toward the Arg186 residue and subsequently positioned the amide nitrogen toward the loop containing Ser92 and Asn93. In structures for all ligands this amide nitrogen was able to form a hydrogen bond with a conserved water molecule in the binding site that bridged interactions with the backbone amide nitrogen from Asn93, with the only exception being one rotamer of **53** (discussed below). Interestingly, reversing this amide connectivity resulted in a 2.5-fold reduction of UCHL3 inhibition when comparing analog **43** to the reverse-amide matched analog **42**. The SAR corroborates the optimal connectivity and placement of the amide to gain two important interactions with the binding site.

Among the three structurally characterized ligands, only the 2-pyridinyl (**43**) and the 3-OH-phenyl (**69**) pendant groups were able to form a new interaction with UCHL3, both to Ser92. In the case of **43**, a water-mediated hydrogen-bond to the side chain of Ser92 (**Figure 5A**). This allowed the 2-pyridine moiety to remain in place and not adopt alternative conformations despite the portion of the molecule being mostly emerged from the pocket and exposed to the solvent (**Figure 5B**). The two water molecules within the active site with **43** were able to help bridge new interactions that account for the improved inhibitory activity compared to the other inhibitors. In the case of **69**, the 3-hydroxy appears to form a direct hydrogen-bond interaction with the backbone amide of Ser92 (**Figure 5G**). Conversely, the methyl and methoxy *meta*-substitutions for analogs **53** and **60** were unable to form new interactions with UCHL3, and both demonstrated conformational flexibility within the binding site in which the pendant group was observed in two different rotamers across the different chains within the asymmetric unit of the same crystal structure. This is illustrated by comparing the pendant phenyl poses for both **53** and **60** between both the Chain A and Chain B structures bound to UCHL3 (**Figure 5C – F**). Interestingly, the placement of the methoxy group for **60** in each rotamer alters the position of the Ser92 side chain, where in the rotamer with the methoxy pointing toward Ser92, the hydroxyl containing side-chain was directed away the ligand (**Figure 5E**). Conversely, when the methoxy was rotated away from Ser92, the hydroxyl side-chain was directed toward the ligand (**Figure 5F**), possibly a result of steric clash. This same rotamer difference was not observed for the slightly smaller 3-methyl substitution of **53** (**Figure 5C** and **D**).

Upon solving the ligand bound structures it was evident that the inhibitor binding position is in an alternate location compared to the natural ubiquitin binding site. This is illustrated with overlaying the structure of UCHL3-**43** with that of UCHL3-ubiqutin (PDB: 1XD3)^31^ in which the inhibitor appears to approach the active site Cys95 from the opposite side of the UCHL3 crossover loop compared to the ubiquitin binding site. This results in conformational flexibility of the crossover loop as in the ubiquitin bound structure the crossover loop is displaced away from ubiquitin to make space for the C-terminal tail to thread into the active site (**Figure 6**, red loop). Conversely, in the ligand-bound structure the ordered part of the cross-over loop is moved more toward the ubiquitin side of UCHL3 to accommodate the ligand on the opposite face (**Figure 6**, green loop). These factors indicate that while the ligand and ubiquitin do not necessarily compete for the same binding site on UCHL3, the dynamics of the cross-over loop placement when either is bound will likely perturb the ability of the other to engage the enzyme. In addition, there will be a clash between Ub’s Gly76 in its C-terminal tail and the chloroacetamide group of the inhibitor, as they both must approach the catalytic cysteine. This will also make the binding mutually exclusive

**Figure 6.**
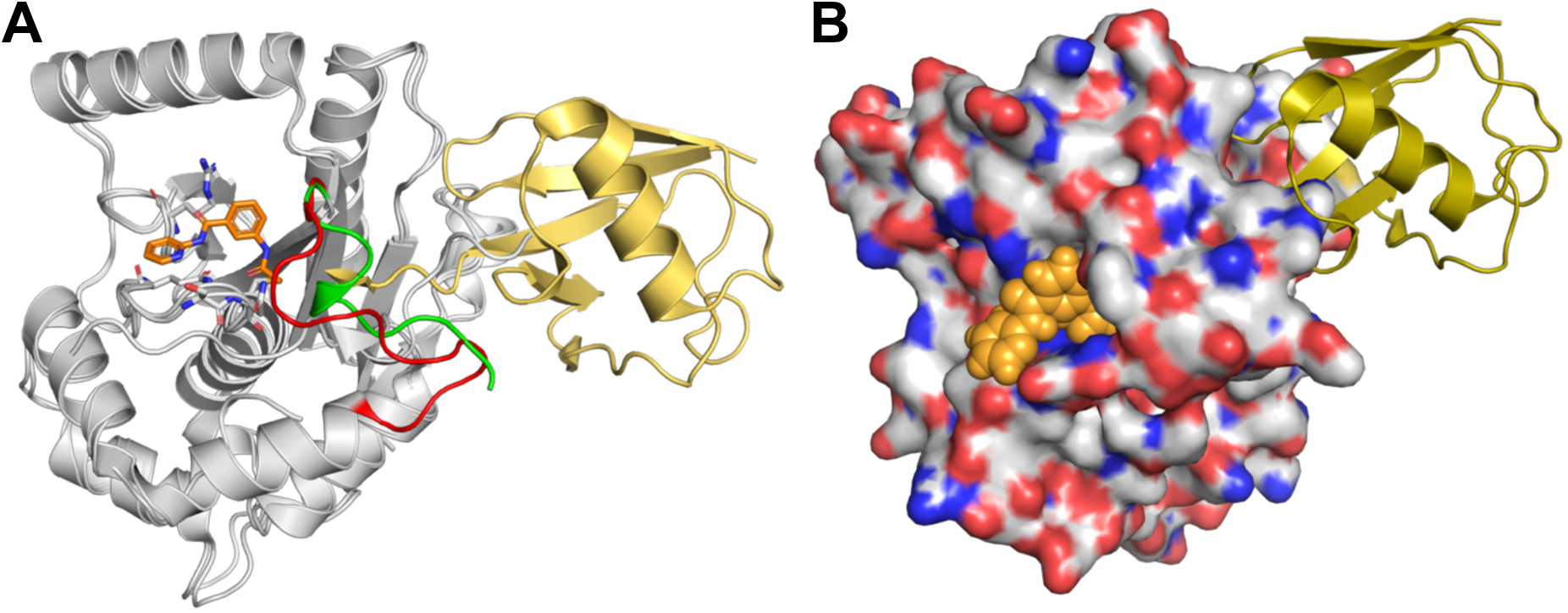
Overlay of UCHL3 ligand bound structure with ubiquitin-bound structure (PDB: 1XD3). (**A**) cartoon representation of UCHL3 (gray) bound to **43** overlaid with UCHL3 bound to ubiquitin (gold cartoon). Cross-over loop of UCHL3 for the ubiquitin bound form shown as red loop. Cross-over loop of UCHL3 for the ligand-bound form shown as green loop. (**B**) Surface representation of UCHL3 showing inhibitor **43** (orange spheres) and ubiquitin (gold cartoon).

### Cellular evaluation of UCHL3 inhibition

To the best of our knowledge, neither of the prior reported UCHL3 inhibitors had been validated for on-target engagement within a cellular context. Therefore, we set out to confirm target engagement for the scaffold in cell lines expressing UCHL3 using a gel-shift assay with HA-Ub-vinyl methyl ester (VME) activity-based probe.^23,32^ First, MiaPaCa2 cells were pre-treated with either **68** or **69** at 0, 5, 10, 20 µM (an additional 30 µM concentration for **69**) for 4 hours. After incubation, cells were lysed and treated with HA-Ub-VME activity-based probe, which reacts with any non-inhibited DUB forming a covalent adduct approximately 8.5 kDa higher in molecular weight than the native DUB. Lysates were then evaluated by immunoblot with α-UCHL3 antibody to visualize molecular weight shift of the HA-Ub-UCHL3 adduct (DMSO control lane) compared to treated samples (**Figure 7**). The data indicates both **68** and **69** successfully inhibit Ub binding to UCHL3 in MiaPaCa2 cells as evidenced by the dose-dependent decrease of HA-Ub-UCHL3 adduct and corresponding increase of unbound UCHL3. Interestingly, analog **68** appears to be more potent in the cellular assay demonstrating nearly full UCHL3 inhibition at 10 µM and fully inhibiting at 20 µM, while **69** was moderately potent at 10 µM with some HA-Ub-UCHL3 adduct remaining at 20 µM (**Figure 7**).

**Figure 7.**
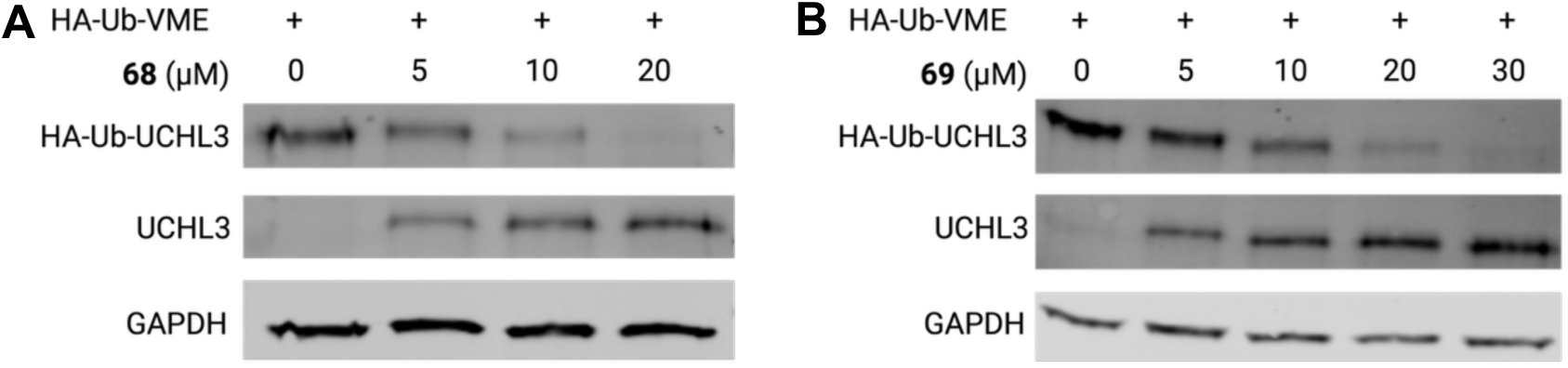
Immunoblot HA-Ub-VME activity-based probe data for MiaPaCa2 cells treated with (**A**) **68** and (**B**) **69**. Blotting for UCHL3 indicated molecular weight shift corresponding to HA-Ub-UCHL3 adduct formation in the control lane (0 µM inhibitor). Increasing dose of molecule blocked formation of the HA-Ub-VME adduct in a dose-dependent manner. GAPDH was used as a loading control.

Next, we evaluated analogs **43** and **69** for target engagement in the same assay using MDA-MB-231 breast cancer cell line (**Figure 8**). The analogs also demonstrated target engagement of UCHL3 in these cells and inhibition of the HA-Ub-VME adduct formation was quantified by normalizing to the DMSO (0 µM) control lanes to have a cellular IC_50_ of approximately 10 µM. The combined data from multiple analogs across two representative cell lines confirm the molecules successfully engaged UCHL3 in a cellular context.

**Figure 8.**
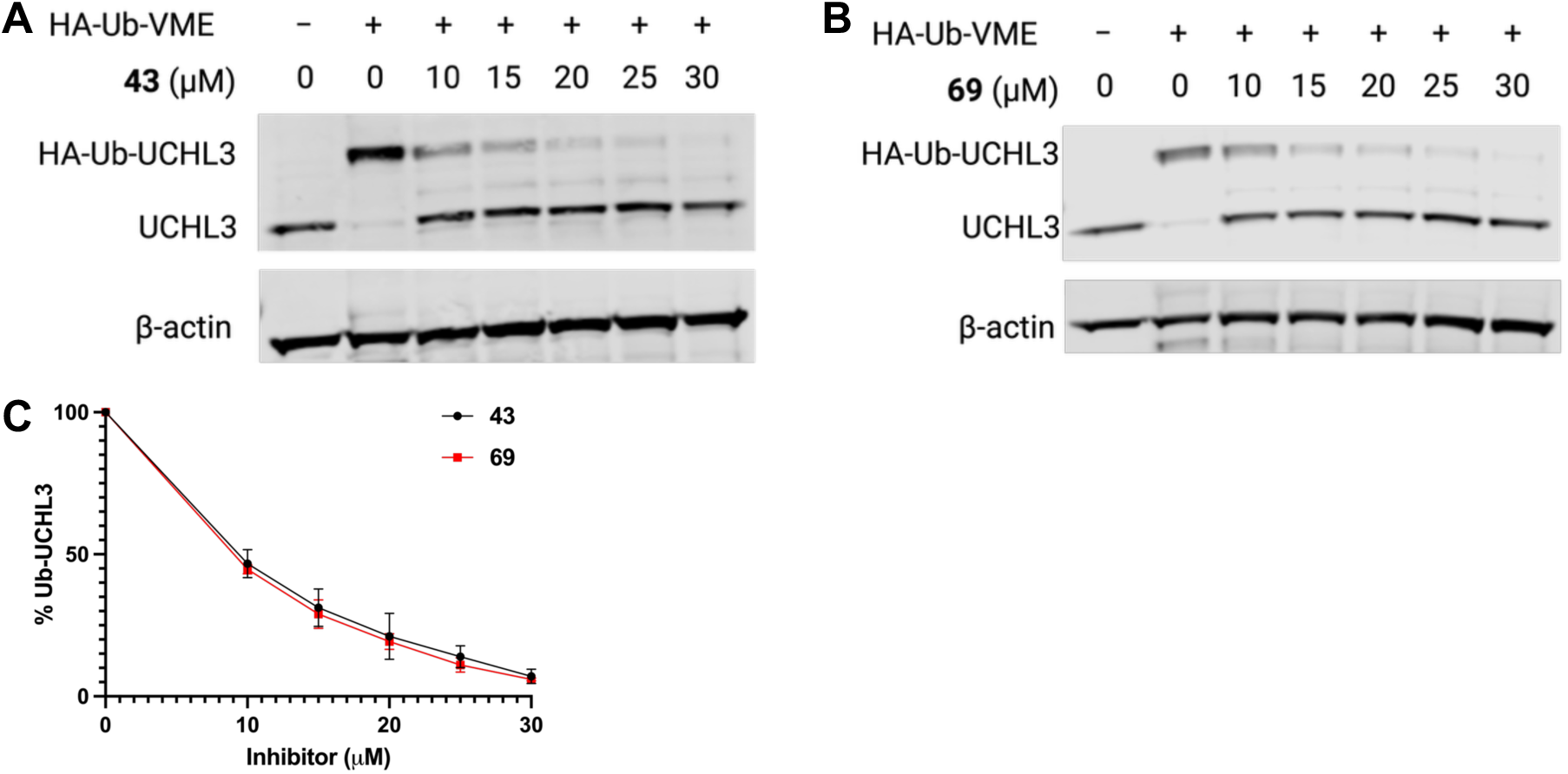
Immunoblot HA-Ub-VME activity-based probe data for MDAMB-231 cells line treated with **(A) 43** and **(B) 69**. Blotting for UCHL3 indicated molecular weight shift corresponding to HA-Ub-UCHL3 adduct formation in the control lane (0 µM inhibitor). Increasing doses of molecule blocked formation of the HA-Ub-VME adduct in a dosedependent manner. b-actin was used as a loading control. (**C**) Quantification of % HA-Ub-UCHL3 normalized to for each inhibitor concentration normalized to 0 µM inhibitor control. Data plotted is mean ± S.D., n = 3 biological reps.

## DISCUSSION

The results present a new irreversible inhibitor chemotype for UCHL3 while reporting both ligand-bound inhibitor crystal structures and intracellular target engagement data. The work offers a broad evaluation of UCHL3 inhibition, complete with SAR and biochemical inhibition corroborated with crystallographic studies and confirmation of cellular target engagement. While there have been two previously reported UCHL3 inhibitors, TCID and perifosine, these prior studies lacked corroboration of biochemical inhibition with cellular target engagement. For example, to the best of our knowledge, there are no definitive target engagement studies using the gold-standard Ub activity-based probe assays for either molecule, and in the case of perifosine, there is a lack of confirmation of *in vitro* biochemical inhibition against recombinant UCHL3. Indeed, in our hands perifosine was unable to inhibit recombinant UCHL3 at concentrations up to 100 µM (data not shown). Thus, there was a gap in the available scientific tool set of small molecules to reliably probe UCHL3.

The findings presented here provide a new series of small molecule covalent inhibitors within the same molecular scaffold that inhibits UCHL3 *in vitro* with clear SAR that is corroborated by structural data and confirms that the molecules inhibit UCHL3 in a cellular setting. Such molecules may be useful for probing the contributions of UCHL3 in different cellular systems and biological contexts and will be best utilized as an orthogonal tool to complement the existing genetic studies involving either knockdown or knockout of UCHL3.

While the findings in this manuscript demonstrate a clear advancement in the field for understanding UCHL3 biology they are not without limitations. Among these limitations is that the scaffold demonstrates modest potency in the low-double digit micromolar range for inhibition against UCHL3 both in biochemical and cellular assays. Additionally, while the scaffold appears to be selective against the closely related recombinant UCHL1 *in vitro*, the broader DUB selectivity and non-specific cysteine labeling still needs to be evaluated. Therefore, the molecules must be utilized in proper context with the dosing concentration range to maximize UCHL3 target engagement and mitigate off-target DUB inhibition to appropriately delineate findings that could be directly related to UCHL3 inhibition.

Future work on this scaffold will include further structure-based optimization to gain potential new interactions with the target. The intracellular selectivity will also be determined at the relevant inhibition concentrations for the top analogs. Further studies on downstream consequences of UCHL3 inhibition with respect to the DUB’s role in cancer biology will also be explored.

## CONCLUSIONS

This interdisciplinary study combined high-throughput screening, medicinal chemistry, biochemical evaluation, and both structural and cellular biology to identify, optimize and biologically evaluate a new irreversible UCHL3 inhibitory scaffold. The top analogs in the study demonstrated IC_50_ values in the 13 – 16 µM range and observed SAR was supported by ligand-bound X-ray crystal structures to UCHL3. The results are complete with characterization of UCHL3 target engagement in two different cell lines using Ub activity-based probe gel shift assays. The results published in this manuscript represent the most comprehensive data from hit-to-probe for UCHL3 in the literature to date and the top analogs **43** and **68**, in terms of UCHL3 biochemical and cellular potency as well as selectivity over UCHL1, will serve as valuable probes to further elucidate the role of UCHL3 in cancer biology.

## EXPERIMENTAL

### Cloning, expression, and purification of UCHL3

The UCHL3 WT gene is amplified with an EcoRI 5’ extension and a XhoI 3’ extension, double digested, and ligated into pGEX-6P-1 (Cytiva). The UCHL1 WT gene was cloned into the ampicillin resistant vector pET-21a containing a C-terminal His_6_ tag. These plasmids were sequence verified and transformed into chemically competent *E. coli* BL21 (DE3) and *E. coli* star DE3 cell respectively, plated onto LB agar containing 100 μg/mL ampicillin, and incubated at 37 °C overnight. Selected colonies were used to inoculate a starter culture containing the same amount of antibiotics and grown at 37 °C for 16 h while shaking. Larger LB media expression cultures containing 100 μg/mL ampicillin were inoculated by a 1:100 dilution of the starter culture and grown at 37 °C while shaking until and OD_600 nm_ = 0.6-0.8. Protein expression was then induced by the addition of 0.5 mM isopropyl β-D-1-thiogalactopyranoside (IPTG) and carried out at 16-18 °C for 18-20 h while shaking. Cells were harvested by centrifugation at 7000 rpm in iterations of 10 min. Cell pellets were flash frozen in liquid nitrogen and stored at -80 °C until further use.

Cell pellets were thawed and resuspended in 1x phosphate buffered saline (PBS) pH 7.0 400 mM KCl. Lysozyme was added to a final concentration of 0.5 mg/mL. Cells were disrupted by high pressure through three passes using a French press. Ultracentrifugation at 75,000 x g for 1 h at 4 °C gave clarified lysate that was applied to either glutathione agarose resin (ThermoFisher Scientific) or Ni-NTA agarose resin (ThermoFisher Scientific) with loading occurring at 4 °C for 1 hr prior to washing. For the GST-tagged UCHL3, the resin was washed with 40-50 CV of 1x PBS 400 mM KCl and then eluted in 5 CV of elution buffer (1x PBS pH 7.4 containing 15 mM reduced glutathione) after another 1 h incubation at 4 °C. Precission protease was added as per the manufacturer’s recommendation and incubated overnight at 4 °C for tag cleavage. The elution was concentrated and buffer exchanged/subject to further purification by size exclusion chromatography by FPLC sporting a HiLoad 16/600 Superdex 75 column (Cytiva) using 1x PBS 1 mM DTT as the mobile phase. Pure fractions as determined by SDS-PAGE Coomassie staining were pooled and concentrated to 20-40 mg/mL, flash frozen in liquid nitrogen and stored at -80 °C until further use.

### UCHL3 WT covalent adduct formation, crystallization and data processing

Adduct formation of UCHL3 with **4**, **43**, **53**, **60**, and **69** was carried out in a 1:5 ratio of enzyme to inhibitor for 2-4 h at room temperature in 1x PBS. Residual DMSO and molecule were removed by buffer exchange via a concentrator and further by size exclusion chromatography as described above. The adduct extent was all verified by intact protein mass spectrometry. The UCHL3-**4** adduct was crystallized by sitting drop vapor diffusion at 21°C. Crystals were grown in one of two conditions: 1) 5% tacsimate pH 8 with 45% PEG8000 or 2) 0.1 M bis-tris propane pH 6.4 2.5 M ammonium sulfate. These crystals diffracted at the Stanford Synchrotron Radiation Lightsource on the 9-2 (λ=0.98) beamline or at the Purdue Center for Macromolecular Crystallography on the home source diffractometer, a Rigaku MicroMax-007 HF rotating anode with confocal VariMax-VHF Arc) Sec optics and a Eiger2 R 4M photon counting detector. Data was processed using HKL3000 in the monoclinic space group P2_1_.

The UCHL3-**43**, UCHL3-**53**, and UCHL3-**60**, and UCHL3-**69** adducts were all crystallized by sitting drop vapor diffusion at 21°C in the same condition of 0.1 M bis-tris propane pH 7 2.5 M ammonium sulfate. Optimization led to varying final condition centered around 100-250 mM bis-tris propane pH 6.1-7.9 containing ammonium sulfate, and 10% glycerol. These crystals were diffracted at the Stanford Synchrotron Radiation Lightsource on the 12-2 (λ=0.98) beamline or the home source Rigaku X-ray MicroMac-007. Data was processed using HKL3000 of CrysAlisPro in the monoclinic space group P2_1_.

### Structural Determination

The inhibitor bound structure of **4** with UCHL3 WT was determined by molecular replacement using the program PHASER in Phenix suite with an input model from the apoUCHL3 WT structure (PDB 1UCH).^33^ Three molecules were observed to be in the asymmetric unit. Ligand coordinates and restraints were generated by using Phenix’s eLBOW and modelled accordingly based on the observed diffraction data. Density accounting the whole molecule was only observed in Chain A. Model building and refinement using the programs COOT and Phenix suite’s phenix refine where the current crystallographic R and R_free_ are 0.21 and 0.25 respectively. Analysis of the Ramachandran plot indicated that 97.13% of residues fell in the most favored regions, 2.55% of residues fell in the allowed regions, and 0.32% were observed in the disallowed region. Electron density is well resolved for the entire molecule and the ligand. A total of 201 water molecules were observed in the asymmetric unit. An average B factor of 51.15 Å^2^ was observed for all atoms in the asymmetric unit.

The inhibitor bound structure of **43**, **53**, **60**, and **69** with UCHL3 WT was determined by molecular replacement using the program PHASER in Phenix suite with an input model from the apoUCHL3 WT structure (PDB 1UCH). Four molecules were observed to be in the asymmetric unit. Ligand coordinates and restraints were generated by using Phenix’s eLBOW and modelled accordingly based on the observed diffraction data. Density accounting the whole molecule was observed in all chains for each molecule. Model building and refinement using the programs COOT and Phenix suite’s phenix refine where the current crystallographic R/R_free_, and other data refinement statistics, are reported in the supporting information.

### IC_50_ Determination

Assays were performed in black 384-well plates (Fisher Scientific Catalog # 12566624) in a final volume of 50 μL. DUB stock solutions were diluted in reaction buffer (25 mM Tris, 50 mM NaCl, 5 mM DTT, 0.1% (w/v) BSA, pH 7.4) to a concentration of 250 pM for UCHL3 or a concentration of 2.5 nM for UCHL1, yielding a final well concentration of 100 pM for UCHL3 or 1 nM for UCHL1. To each well, 10 μL of 5× small molecules in reaction buffer (500, 250, 125, 62.5, 31.25, 15.63, 7.82, 3.91, 1.95, 0 μM) and 20 μL of UCHL3 or UCHL1 solution were added. This was incubated at room temperature for 1 h. Reactions were initiated by adding 20 μL of 500 nM Ub-Rho for UCHL3 or 250 nM for UCHL1 (Boston Biochem U-555, final concentration in the well: 200 nM for UCHL3, 100 nM for UCHL1) and were read immediately for 20 min on a Synergy Neo 2. IC_50_ values were calculated using GraphPad Prism 10, and the standard deviation was calculated over three independent experiments.

### Preincubation time-dependent kinetic determination

Assays were performed in black 384-well plates (Fisher Scientific Catalog # 12566624) in a final volume of 50 μL. UCHL3 was diluted in reaction buffer (25 mM Tris, 50 mM NaCl, 5 mM DTT, 0.1% (w/v) BSA, pH 7.4) to 62.5 pM, yielding a final well concentration of 25 pM. To each well, 10 μL of 5× small molecules in reaction buffer (500, 250, 125, 62.5, 31.25, 15.63, 7.82, 3.91, 1.95, 0 μM) and 20 μL of UCHL3 or UCHL1 solution were added. This was incubated at room temperature for different time periods (0, 5, 10, 15, 30, 60, 90, 120 min) to obtain time-dependent results. Reactions were initiated by adding 20 μL of 500 nM Ub-Rho (Boston Biochem U-555, final concentration in the well: 200 nM) and were read immediately for 2 hr on a Synergy Neo 2. The % activity was plotted vs. preincubation time at each concentration in Prism 10 to obtain *k*_obs_ for inhibition using a one-phase decay model. The *k*_obs_ were plotted as a function of inhibitor concentration and then fit to the equation Y = *k*_inact_*X/(*K*_I_+X) to obtain *k*_inact_, *K*_I_, and *k*_inact_/*K*_I_ for compound **68**. The *k*_obs_ were graphed vs. inhibitor concentration, and the slope of the linear fit was determined to be *k*_inact_/*K*_I_ for compounds **43** and **69**.

### Cellular Target Engagement Assay using HA-Ub-VME activity-based probe

This assay was run according to same protocol reported previously.^23^ Briefly, MiaPaCa2 or MDA-MB-231-luc3 pLKO cells were treated with the indicated concentrations of inhibitor and incubated for 4h at 37 °C, following which the cells were washed, scraped and stored at -80 °C. Cell pellets were resuspended in the lysis buffer (50 mM Tris pH 7.6, 150 mM NaCl, 5 mM MgCl2, 0.5 mM EDTA, 0.5% NP-40, 10% v/v glycerol with freshly added protease inhibitors) and incubated on ice for 30 min with vortexing every 10 min. Cell lysate was clarified by centrifugation at 21,000 × g for 30 min, and the supernatant was collected. Total protein concentration was determined using BCA assay with BSA as standards, according to manufacturer’s protocol and whole cell extracts were made to1 µg/µL. For the DUB assay, 30 µl extract was used for each reaction and DTT and ATP were added fresh to a final concentration of 5 mM, followed by 0.5 µM HA-Ub-VME. This was incubated at RT for 20 min before quenching with 4× LDS sample buffer. Samples were run on a 4-12 % bis-tris gradient gel followed by immunoblotting. The following primary antibodies were used: anti-UCHL3 (ab126621, Abcam), and anti-Actin (sc-47778, Santa Cruz). Fluorescent secondary antibodies (Licor IRDye 680RD goat anti-mouse and Licor IRDye 800CW goat anti-rabbit) were used. Images were collected on a Licor Odyssey system. Densitometry measurements were done using Licor ImageStudio software.

## Acknowledgments

The authors gratefully acknowledge the support for both shared resources and pilot grant funds from the Purdue Institute for Cancer Research NIH grant P30 CA023168, R01GM126296 (C.D.) and the NIH Ruth L. Kirschstein NRSA (F31CA275390 to R.P.). The authors also gratefully acknowledge Purdue Institute for Drug Discovery (PIDD) for both high-throughput screening and programmatic grants for funds. Acknowledgement is also given to the Chemical Genomics Facility, a core facility of PIDD and NIH-funded Indiana Clinical and Translational Sciences Institute for helping conduct the HTS. Finally, the authors would like to acknowledge the assistance of the Purdue Center for Macromolecular Crystallization core facility for use of equipment and resources to identify crystallization conditions and NIH grant S10 OD030507 for home X-ray source equipment. Figures were created using Biorender.com.

## Author Contributions

The manuscript was written through contributions from all authors. All authors gave approval of the final version of the manuscript. Conceptualization: C.D., D.P.F. High-throughput screening and analysis: R.P., N.P., R.D.I., C.D., D.P.F. Molecule synthesis: N.C.B., M.L. H.M., R.D.I. Biochemical evaluation: M.L., N.P., R.P. Crystallographic evaluation: R.P., N.P., C.D. Cellular evaluation: A.D., B.N.H., E.G.S, M.B.B., B.L.A-P. M.K.W. Writing Initial drafts: N.C.B., M.L. D.P.F. Editing and refinement of final draft: All.

## Data availability

Small molecule characterization data will be provided upon request. Small molecule dry powders will be provided upon request pending sufficient stock.

UCHL3 inhibitor bound structure coordinates and processed diffraction data have been deposited in the protein data bank under the accession codes: pdb_000013CU, pdb_000013CV, pdb_000013CW, pdb_000013DF, pdb_000035YH.

