## Supporting Information File for "Identification, optimization, and structural elucidation of chloroacetamide scaffold as covalent inhibitors for Ubiquitin C-terminal Hydrolase L3"

| **Table S1. Crystallography statistics and refinement for analogs 4 and 43**. | | |
|---|---|---|
| | UCHL3-**4** | UCHL-**43** |
| | PDB: pbd_000013CU | PDB: pbd_000013CV |
| | Data Collection | |
| Beamline | SSRL BL 9-2 | SSRL BL 12-2 |
| Wavelength (Å) | 0.98 | 0.98 |
| Resolution range (Å) | 40.53-1.95 (2.02-1.95) | 40.53-1.95 (2.02-1.95) |
| Space group | P 1 21 1 | P 1 21 1 |
| Cell dimensions | | |
| a, b, c (Å) | 44.5, 81.1, 100.7 | 48.029, 85.605, 124.078 |
| α, β, γ (˚) | 90, 98.6, 90 | 90, 96.8, 90 |
| Total reflections | 97851 (9772) | 138078 (12608) |
| Unique reflections | 49972 (4349) | 71473 (6498) |
| Redundancy | 6.2 (6.2) | 2.9 (2.7) |
| Completeness (%) | 96.9 (83.7) | 97.39 (88.96) |
| $R_{meas}$ | 0.162 (2.021) | 0.141 (1.34) |
| $R_{pim}$ | 0.064 (0.792) | 0.058 (0.56) |
| I/σ(I) | 11.825 (1.24) | 10.87 (1.61) |
| CC1/2 | 0.993 (0.573) | 0.995 (0.592) |
| Wilson B ($Å^2$) | 37.53 | 33.95 |
| | Refinement | |
| Copies/A.S.U. | 3 | 4 |
| Resolution (Å) | 1.95 | 1.95 |
| Rwork / Rfree | 0.20 / 0.25 | 0.20 / 0.24 |
| No. nonhydrogen atoms | 5231 | 7209 |
| Protein | 4994 | 6778 |
| Ligand | 62 | 174 |
| Water | 201 | 305 |
| B factor ($Å^2$) | 51.15 | 50.23 |
| Protein | 51.38 | 50.22 |
| Ligand | 50.20 | 60.56 |
| Water | 45.64 | 46.06 |
| R.m.s.d | | |
| Bond lengths | 0.023 | 0.028 |
| Bond angles | 1.22 | 2.03 |
| Ramachandran (favored/allowed/outliers) | 97.13/2.55/0.32 | 97.16/2.37/0.47 |
| Clash Score | 15.09 | 20.32 |

| **Table S2. Crystallography statistics and refinement for analogs 53 and 60**. | | |
|---|---|---|
| | UCHL3-**53** | UCHL-**60** |
| | PDB: pbd_000013CW | PDB: pbd_000013DF |
| | Data Collection | |
| Beamline | SSRL BL 12-2 | SSRL BL 12-2 |
| Wavelength (Å) | 0.98 | 0.98 |
| Resolution range (Å) | 47.4-2.48 (2.57-2.48) | 42.68-1.84 (1.90-1.84) |
| Space group | P 1 21 1 | P 1 21 1 |
| Cell dimensions | | |
| a, b, c (Å) | 47.814, 85.406, 124.129 | 48.202, 85.305, 124.367 |
| α, β, γ (˚) | 90, 97.7, 90 | 90, 97.6, 90 |
| Total reflections | 67199 (6570) | 164334 (15269) |
| Unique reflections | 34407 (3343) | 84042 (7734) |
| Redundancy | 2.0 (2.0) | 2.0 (2.0) |
| Completeness (%) | 97.61 (95.73) | 97.07 (90.01) |
| $R_{meas}$ | 0.074 (0.825) | 0.067 (0.968) |
| $R_{pim}$ | 0.052 (0.582) | 0.047 (0.685) |
| I/σ(I) | 8.65 (1.66) | 8.81 (1.23) |
| CC1/2 | 0.997 (0.525) | 0.994 (0.463) |
| Wilson B (Å$^2$) | 52.22 | 34.14 |
| | Refinement | |
| Copies/A.S.U. | 4 | 4 |
| Resolution (Å) | 2.48 | 1.84 |
| Rwork / Rfree | 0.21 / 0.27 | 0.19 / 0.23 |
| No. nonhydrogen atoms | 6757 | 7356 |
| Protein | 6568 | 6835 |
| Ligand | 155 | 159 |
| Water | 94 | 402 |
| B factor (Å$^2$) | 68.24 | 48.60 |
| Protein | 68.42 | 48.67 |
| Ligand | 67.09 | 45.24 |
| Water | 56.88 | 48.31 |
| R.m.s.d | | |
| Bond lengths | 0.005 | 0.135 |
| Bond angles | 1.03 | 1.34 |
| Ramachandran (favored/allowed/outliers) | 97.36/2.52/0.12 | 97.30/2.58/0.12 |
| Clash Score | 23.63 | 13.07 |

| **Table S3. Crystallography statistics and refinement for analog 69.** | |
|---|---|
| | UCHL3-**69** |
| | PDB: pbd_000035YH |
| | Data Collection |
| X-ray source | Rigaku X-ray MicroMax-007 |
| Detector | Eiger2 R 4M |
| Wavelength (Å) | 1.54 |
| Resolution range (Å) | 20.15 – 2.6 (2.693-2.6) |
| Space group | P 1 21 1 |
| Cell dimensions | |
| a, b, c (Å) | 47.87, 84.99, 123.86 |
| α, β, γ (˚) | 90, 97.9, 90 |
| Total reflections | 121247 |
| Unique reflections | 30188 (3032) |
| Redundancy | 4.9 (5.0) |
| Completeness (%) | 99.02 (99.21) |
| I/σ(I) | 0.9 |
| CC1/2 | 0.37 |
| Wilson B (Å$^2$) | 56.35 |
| | Refinement |
| Copies/A.S.U. | 4 |
| Resolution (Å) | 2.60 |
| Rwork / Rfree | 0.25 / 0.31 |
| No. nonhydrogen atoms | 6222 |
| Protein | 6115 |
| Ligand | 132 |
| Water | 27 |
| B factor (Å$^2$) | 71.68 |
| Protein | 71.83 |
| Ligand | 65.55 |
| Water | 58.19 |
| R.m.s.d | |
| Bond lengths | 0.026 |
| Bond angles | 0.66 |
| Ramachandran (favored/allowed/outliers) | 95.69/4.05/0.26 |
| Clash Score | 29.38 |

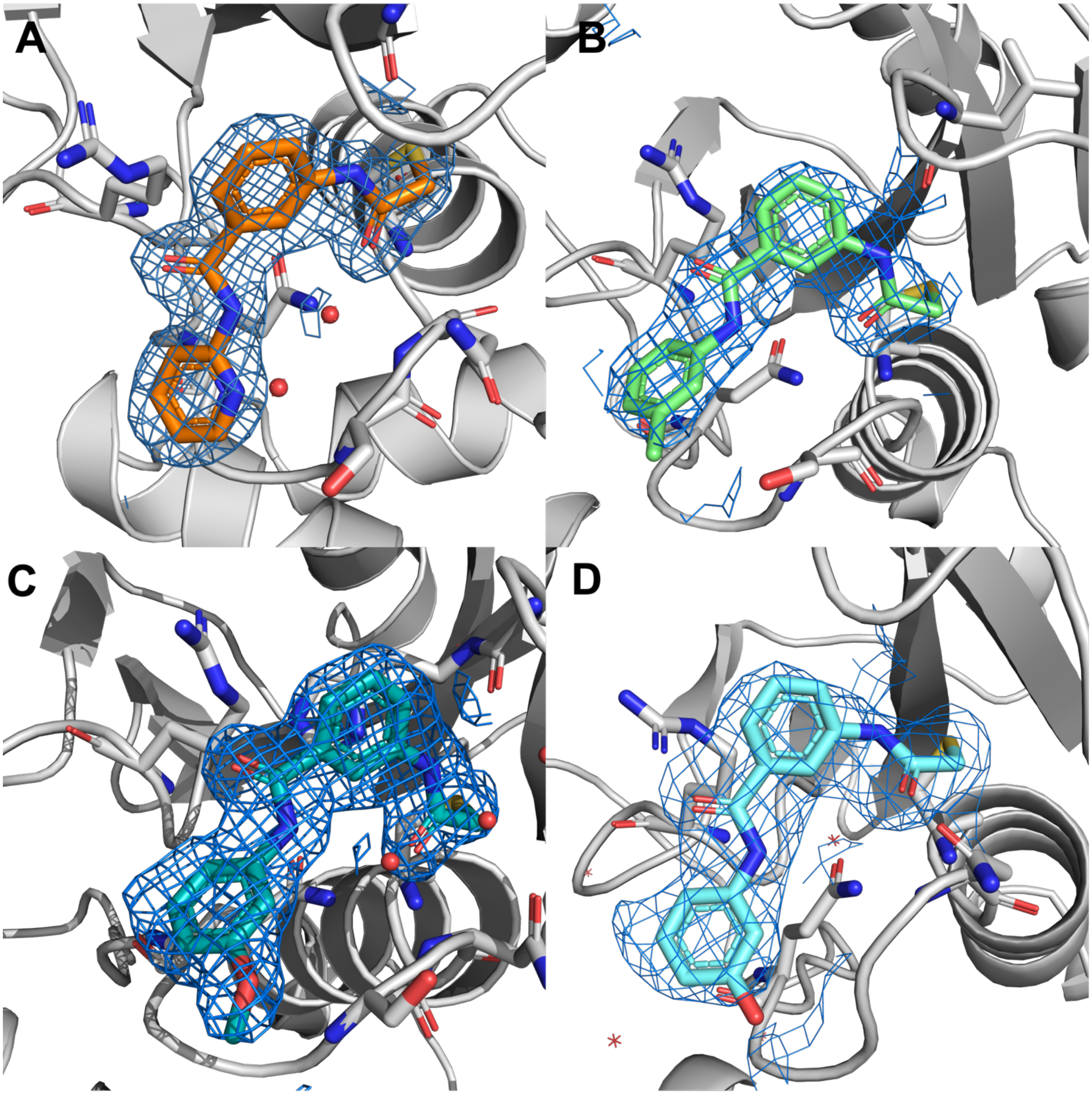

**Figure S1.** Omit $|2F_0 - F_c|$ electron density maps for UCHL3 in complex with analogs. (**A**) **43** (orange sticks) with UCHL3 Chain A (gray). (**B**) **53** (mint sticks) with UCHL3 Chain A (gray). (**C**) **60** (teal sticks) with UCHL3 Chain A (gray). (**D**) **69** (sky blue sticks) with UCHL3 Chain A (gray).
